# High-Quality Genome Assembly and Annotation of the California Harvester Ant *Pogonomyrmex californicus* (Buckley, 1867)

**DOI:** 10.1101/2020.09.01.277236

**Authors:** Jonas Bohn, Reza Halabian, Lukas Schrader, Victoria Shabardina, Raphael Steffen, Yutaka Suzuki, Ulrich R. Ernst, Jürgen R. Gadau, Wojciech Makałowski

## Abstract

The harvester ant genus *Pogonomyrmex* is endemic to arid and semiarid habitats and deserts of North and South America and California harvester ant *Pogonomyrmex californicus* is the most widely distributed *Pogonomyrmex* species in the North America. *P. californicus* colonies are usually monogynous, i.e. a colony has one queen. However, in a few populations in California, primary polygyny evolved, i.e. several queens cooperate in colony founding after their mating flights and continue to coexist in mature colonies. Here, we present high quality genome assembly and annotation of *P. californicus*. The size of the assembly is 241 Mb, which is in good agreement with previously estimated genome size and we were able to annotate 17,889 genes in total, including 15,688 protein-coding ones with BUSCO completeness at the 95% level. This high quality genome will pave the way for investigations of the genomic underpinnings of social polymorphism in queen number, regulation of aggression, and the evolution of adaptations to dry habitats in *P. californicus*.

---

Ants (Hymenoptera: Formicidae) are important components of almost all terrestrial ecosystems and more than 16,000 species have been described so far (AntWeb, version 8.41, California Academy of Science, online at https://www.antweb.org; accessed on August 19, 2020). The majority of them, over 6,900, belong to the highly diverse subfamily Myrmicinae ants (AntWeb, version 8.41, California Academy of Science, online at https://www.antweb.org; accessed on August 19, 2020). Currently, 40 assembled ant genomes are available at NCBI (Entrez “Genome” accessed on August 19, 2020).

The harvester ant genus *Pogonomyrmex* is endemic to arid and semiarid habitats and deserts of North and South America (Buckley 1867; Cole 1968; Snelling et al. 2009). This genus thrives in extremely dry habitats, e.g. Death Valley, Anza Borega and evolved seed harvesting behavior independently from the Old World harvester ant genus *Messor*. Members of the genus *Pogonomyrmex* are a very conspicuous element of the deserts in the Southwest of the USA and have been studied extensively (De Vita 1979; Rissing et al. 2000; Lighton and Turner 2004; Clark and Fewell 2014; Helmkampf et al. 2016; Overson et al. 2016). Within this genus several interesting traits like social parasitism, genetic caste determination and social polymorphism in terms of queen number have evolved (Cole 1968; Rissing et al. 2000; Julian et al. 2002). Arguably, the most widely distributed *Pogonomyrmex* species in the Northern America is *P. californicus* (Johnson 2002). *P. californicus* colonies are usually monogynous, i.e. a colony has one queen. However, in a few populations in California, primary polygyny evolved, i.e. several queens cooperate in colony founding after their mating flights and continue to coexist in mature colonies (Rissing et al. 2000; Johnson 2004; Shaffer et al. 2016). The Red Imported Fire Ant, *Solenopsis invicta* and several other *Formica* ant species have a similar social polymorphism, which has been shown to be due to a supergene (Wang et al. 2013; Yan et al. 2020). This discovery was only possible by Nextgen Sequencing and the availability of genomic information for these species. Of approximately seventy described *Pogonomyrmex* species only *P. barbatus* (AntWeb, version 8.41, California Academy of Science, online at https://www.antweb.org; accessed on August 19, 2020) has its genome sequenced, assembled and annotated (Smith et al. 2011). Five other species of this genus (*P. anergismus, P. colei, P. imberbiculus, P. occidentalis*, and *P. rugosus*) have nuclear genomes partially sequenced but none has so far been processed and only raw reads are available at NCBI’s Sequence Reads Archive (SRA). Sequences *of P. rugosus, P. anergism* and *P. colei* have been aligned to the *P. barbatus* genome for a study of gene gains/losses in social parasitic ants (Smith et al. 2015).

The genome sequencing and annotation of *P. californicu*s will result in a better understanding of genomic sequence and structural variations and evolution in Formicidae in general and in the myrmicine genus *Pogonomyrmex* in particular. It will also pave the way for investigations of the genomic underpinnings of social polymorphism in queen number, regulation of aggression, and the evolution of adaptations to dry habitats in *P. californicus*.

## MATERIALS AND METHODS

### Samples and transcriptome data source

The nuclear DNA was extracted from thirteen haploid males from a single polygynous colony collected 2016 from Pine Valley, California, USA (32.819761, -116.521512; N32 49 11 – W116 31 17). Previously published transcriptome data (Helmkampf et al. 2016) were downloaded from NCBI’s Sequence Read Archive (BioProject accession number PRJDB4319). In addition, Oxford Nanopore sequencing (MinION) was performed on RNA extracted from workers of five polygynous colonies also collected in 2016 from Pine Valley, California, USA. These reads are accessible at NCBI with BioProject accession number PRJNA622899.

### Genome Sequencing and Assembling

DNA from thirteen male ants was isolated using Qiagen MagAttract HMW DNA Kit with the protocol for tissue DNA extraction according to the protocol from October 2017. It resulted in 4,575 ng of DNA of which 1.2 ng was used for 10x Genomics Chromium sequencing approach. The sequencing library was prepared according to Chromium™ Genome Library Kit standard protocol (Manual Part Number CG00022) and Illumina HiSeq 3000 system was used to sequence the library. The quality of the produced reads was checked using FastQC software, version 0.11.5 (Andrews 2010). We performed neither filtering nor trimming on these linked reads to avoid losing any information. We used the *de novo* assembler Supernova, version 2.1 (Weisenfeld et al. 2017) with following parameters: --maxreads=156111200 and -- accept-extreme-coverage. Maximum number of reads was set to 75x effective coverage of the genome, which was chosen based on a set of Supernova runs with different coverages to obtain optimal parameters as a trade-off between genome size, BUSCO assessment (see below) and N50 coverage (see Figure 1) and assuming *P. californicus* genome size 244.5 Mb (http://www.genomesize.com). Subsequently, the resulting genome assembly was polished by three rounds of Pilon, version 1.23, processing (Walker et al. 2014). For this step 678,988,626 one-hundred bp reads from an independent Illumina sequencing were added to the 269,953,173 linked-reads used for the genome assembly bringing the number of reads used in the polishing step to almost one billion. For this additional sequencing standard Illumina protocol was used to prepare a sequencing library, which was sequenced using Illumina HiSeq 3000 system.

**Figure 1:**
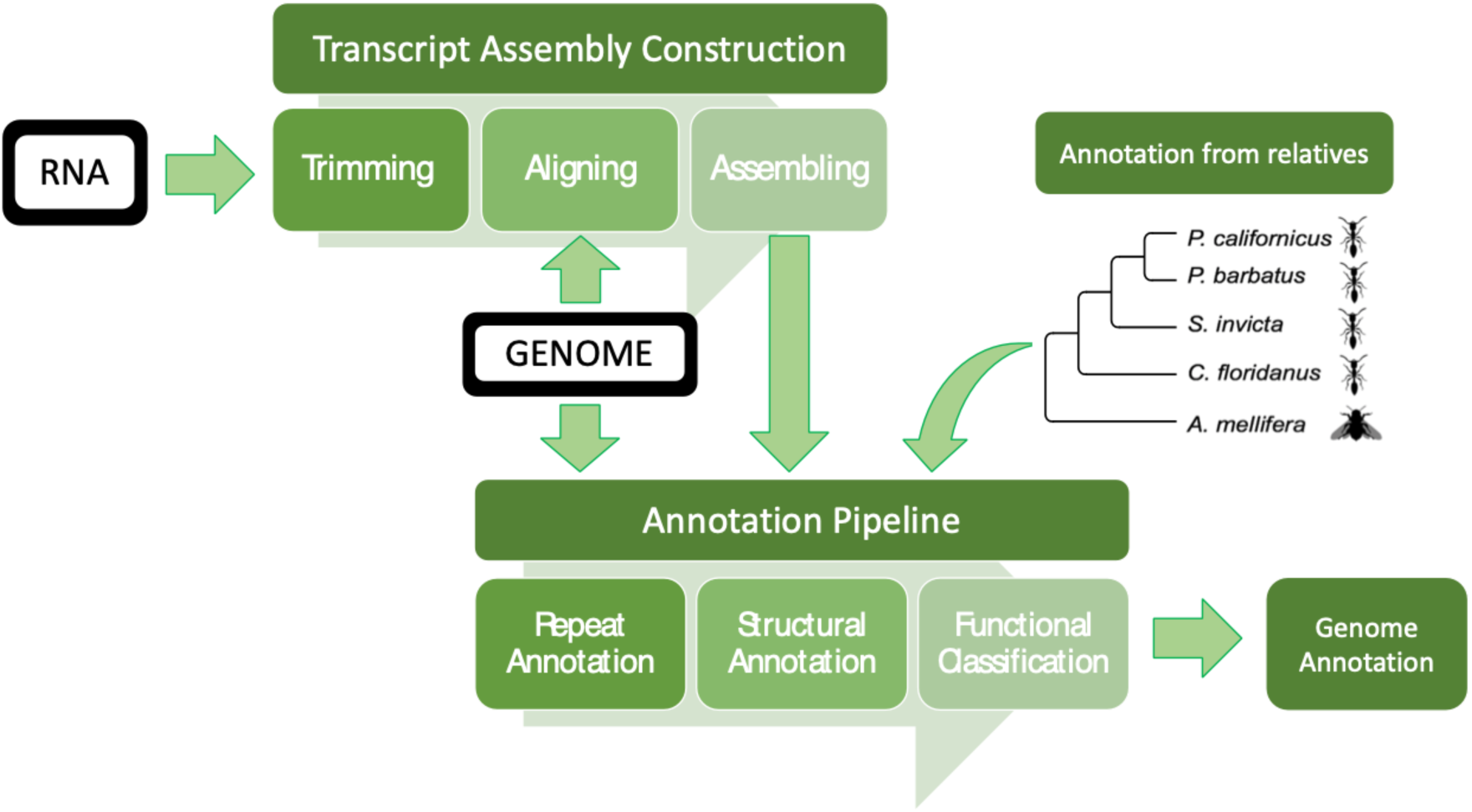
Overview of the annotation workflow. The workflow includes the construction of the transcript assembly (upper part) and the pipeline for the genome annotation (lower part). The transcript assembly and annotations from related species are providing evidence for the annotation of protein-coding genes.

### Transcript Sequencing and Assembling

For transcriptome analysis, MinION long read RNA sequencing of the entire body of ant workers from a laboratory colony (pleometrotic colony from Pine Valley, USA) was performed. We extracted RNA using Monarch® Total RNA Miniprep Kit (New England BioLabs GmbH, Frankfurt, D, E2010). Material was ground in Mixer Mill 200 (Retsch GmbH, Haan, D) in protection reagent. Quality check was performed with Bioanalyser, Nanophotometer and Qubit. The library was prepared from 5 µg of total RNA using cDNA-PCR sequencing kit SQK_PCS_9035_v108_revD_26.6.17 (Oxford Nanopore Technologies, Oxford, UK). The library was sequenced using a MinION and a flow cell FLO-MIN107 R9 (Oxford Nanopore Technologies, Oxford, UK). ONT’s albacor software, version 2.3.1 with standard parameters was used for base calling and only sequences that passed standard quality check (placed in “pass” folder by basecaller) were used for further analyses.

The RNA NGS Illumina reads from Helmkampf et al. (2016) were aligned employing Hisat, version 2.1.0 (Kim et al. 2015) for a genome dependent assembly. A genome independent transcript assembly was done using Trinity version 2.8.4 (Grabherr et al. 2011) on the NGS RNA-Seq data and using the Trinity assembly provided by Helmkampf et al. (2016). In addition, minimap2 version 2.17 within FLAIR pipeline version 1.4 was used for aligning nanopore long reads and the Trinity assemblies to the genome. Finally StringTie2 version 2.0.1 (Kovaka et al. 2019) was employed in order to link the different transcript assemblies filtered by a minimum FPKM of 0.14 as it was also done by Helmkampf et al. (2016).

### Repeat Annotation

We used two independent pipelines for *de novo* repeats discovery, namely RepeatModeler, version 1.0.11 (http://repeatmasker.org/RepeatModeler/) and REPET, version 2.5 (Flutre et al. 2011). The obtained libraries were merged with Hymenoptera specific repeats from RepBase, version 22.07 (Bao et al. 2015). TEclass software, version 2.1.3 (Abrusán et al. 2009) was used for classification of consensus sequences lacking TE-family assignment. Finally, we removed sequences sharing more than 90% identity by employing CD-HIT version 4.7 (Fu et al. 2012). This resulted in the library consisting of 2,595 consensus sequences, which was used to annotate repeats in *P. californicus* genome using RepeatMasker, version 4.0.7 (Smit et al. 2013).

### Protein-coding Gene Annotation

The identification of protein-coding genes in the newly assembled genome of *P. californicus* was carried out by GeneModelMapper (GeMoMa) version 1.6.1 (Keilwagen et al. 2018) followed by MAKER2, version 2.31.10 (Holt and Yandell 2011) (see Figure 1). We used annotation of four insect species (*P. barbatus, Solenopsis invicta, Camponotus floridanus*, and *Apis mellifera*) to run GeMoMa. These annotations were downloaded from NCBI (see Supplementary table 1). GeMoMa was run for each reference species separately and the results were merged using GeMoMa annotation filter (GAF). Next, four runs of MAKER2 was used to refine genome annotation. MAKER2 was used with the following data: GeMoMa predictions, transcript assembly, transcript and protein annotations from relative species, and RepeatMasker annotation (see above). AUGUSTUS (Stanke and Morgenstern 2005), which is a part of MAKER2 pipeline, was trained on AUGUSTUS reference model from *Nasonia* for the first run and trained on the created *P. californicus* reference model by applying BUSCO version 3.0.2 (Waterhouse et al. 2018) for the third run. In addition, SNAP (Korf 2004) was performed for the last three MAKER2 run and trained on Hidden Markov Model (HMM) reference models from gene predictions of the previous run with minimum length of 50 amino acids and maximum annotation edit distance (AED) of 0.25. Redundant identical transcripts and proteins within the final predictions of MAKER2 were filtered with CD-Hit, version 4.7 (Fu et al. 2012).

### Functional Classification of Protein-Coding Genes

The functional classification of the unique protein-coding genes (PCGs) were based on sequence similarity. NCBI’s non-redundant (nr) protein database was searched using BLASTP, version 2.2.31 (Altschul et al. 1990) with default settings except e-value set to 1e-6 and coverage threshold as described below. We considered three possibilities for query and reference sequences overlap. First, the exact matches of the BLAST alignment covers more than 70% to the reference protein and the query protein. In this case the query protein is similar to the reference protein. Second, if the query protein is just a part of the reference protein, the BLAST matches will cover more that 70% of the query sequence but less than 70% of the reference sequence. Lastly, the reference sequence might be included in the query protein. In this case, the BLAST matches are covering more than 70% of the reference protein but less than 70% of the query protein. This allowed adding the functional description from the reference protein (annotated protein in the nr database) to the protein query (*P. californicus* protein predicted by MAKER2 annotation). Further downstream analysis was done with Interproscan version 5.30 (Jones et al. 2014) for deletion of protein domain residues in classified and non-classified proteins. This analysis includes several pipelines including PANTHER, Pfam, Gene3D, SUPERFAMILY, MobiDBLite, ProSiteProfiles, SMART, CDD, Coils, PRINTS, TIGRFAM, PIRSF, Hamap, ProDom, and SFLD.

### Odorant Receptor (OR) Annotation

We annotated odorant receptors (ORs) for the genomes of *P. californicus* and *P. barbatus* using manually curated OR gene models from three other ant species, *Acromyrmex echinator, Atta cephalotes* and *Solenopsis invicta* (McKenzie et al. 2016). Initial OR gene models were annotated with exonerate (version 2.4.0) and GeMoMa, version 1.4, and combined with Evidence Modeler, version 1.1.1 (Haas et al. 2008). All models were screened for the 7tm_6 protein domain typical for insect odorant receptor proteins with PfamScan version 1.5. All genes were further assigned to different OR protein subfamilies by aligning the protein sequence against a set of OR subfamily reference sequences (Sean McKenzie, pers. comm.).

Protein alignment was calculated with MAFFT (Katoh et al. 2002) using following parameters: -globalpair=T -keeporder=T, -maxiterate=16. The resulting alignment was trimmed employing trimal with the parameters: -keepheader=T -strictall=T) (Capella-Gutiérrez et al. 2009). The phylogenetic tree of all predicted OR gene models in both ant species was inferred with FastTreeMP (Price et al. 2010) with following settings: -pseudo -lg -gamma.

### Annotation of non-protein-coding genes

In addition to the protein-coding genes, non-coding genes were annotated as well. Genes for tRNAs have been annotated with tRNAscan-SE, version 2.0.3 (Chan et al. 2019). Other types of ncRNAs including rRNAs, snRNAs, snoRNAs, miRNAs, and lncRNAs have been predicted by Infernal, version 1.1.2 (Nawrocki and Eddy 2013). To this end, we downloaded the Rfam library (release 14.1) of covariance models along with the Rfam clan file (https://rfam.xfam.org). Afterwards, cmscan, a built-in Infernal program, was used to annotate the RNAs represented in the Rfam library in the genome under study. Eventually, the lower-scoring overlaps were removed and the final results were used to generate the gff file containing the annotation of non-coding RNA genes. In addition, we searched for homologues of lncRNA genes from *P. barbatus* (based on annotation of assembly from Supplementary table 1) in *P. californicus* using Splign, version 2.1.0 (Kapustin et al. 2008). Genes where the exons detected by Splign cover more than 90% of *P. barbatus* lncRNA genes were classified as lncRNA genes in the *P. californicus* genome assembly.

### Comparative genomic analysis

The LAST aligner version 909 was used for whole-genome alignments (Kielbasa et al. 2011). The *P. califonicus* and *P. barbatus* genome assemblies were aligned in order to find cognate genes and to search for conserved synteny. We used BEDTools intersect, version 2.27.1 (Quinlan and Hall 2010) to compare the genome annotations and estimate the proportion of shared genes.

### Assembly and annotation quality assessment

We assessed the quality of our assembly and annotation using BUSCO, version 3.0.2 (Waterhouse et al. 2018) and DOGMA web server (Dohmen et al. 2016; Kemena et al. 2019). For BUSCO analyses, we used the Hymenoptera specific single-copy orthologous genes from OrthoDB version 9 (Zdobnov et al. 2017). For DOGMA, we employed the insect domain core set from Pfam, version 32.

## Data Availability

All analyses, including the assembly and the annotation pipeline are available under http://www.bioinformatics.uni-muenster.de/publication_data/P.californicus_annotation/index.hbi. The raw sequencing data are available at NCBI Sequence Read Archive under accession number PRJNA622899 (https://www.ncbi.nlm.nih.gov/sra/?term=PRJNA622899).

## RESULTS AND DISCUSSION

### Sequencing results

We performed two rounds of NGS genomic sequencing and transcriptome sequencing using nanopore long reads technology. We obtained 339,494,313 of 100 bp pair-end reads after standard Illumina sequencing and 269,953,173 of 150 bp linked-reads using 10x Genomics approach. Only the latter reads were used for the genome assembly. Additionally, MinION transcriptome sequencing resulted in 394,085 reads ranging between 49 and 6,182 bp. N50 of the set was 660 bp and total size 241.6 Mbp with median read quality score of 7.8.

### Genome assembly and evaluation

We assembled a high quality draft *P. californicus* genome using linked-read 10x Genomics approach and the Supernova assembler. Assuming 244.5 Mb as a genome size of *P. californicus* (http://www.genomesize.com) our 10X Genomics data coverage was 162x. Supernova was originally designed for *de novo* assemblies of human genomes (Weisenfeld et al. 2017). However, recently it has been successfully used for non-human genome assemblies (Ozerov et al. 2018; Wang et al. 2019; Lu et al. 2020). For human genomes, a 56x coverage is recommended, however, since there is not much information on the optimal coverage for non-model genomes, we performed a series of assemblies. We re-sampled our sequencing data to obtain coverages ranging from 47x to 162x (see Figure 2). In order to minimize the number of artificially duplicated and missing BUSCO genes, we decided that a coverage of 75x is optimal for the assembly of the *P. californicus* genome (see Figure 2). Based on this assembly (75x coverage), the *P. californicus* draft genome consisted of 6,793 contigs totaling in 240,287,203 bp with about 13 undetermined nucleotides (Ns) per 1 kb. The Supernova assembly was followed by three rounds of polishing by Pilon. This resulted in further improvement of the assembly, with the final assembly of 241,081,918 bp and reduced number of N characters (see Table 1).

**Table 1.** Improving of the *P. californicus* assembly by polishing procedure.

|  | Assembly size (bp) | Number of Ns per 1 kb |
| --- | --- | --- |
| **Draft assembly** | 240,287,203 | 13.00 |
| **1st round of polishing** | 240,787,628 | 11.91 |
| **2nd round of polishing** | 241,014,132 | 11.64 |
| **3rd round of polishing** | 241,081,918 | 11.53 |

**Figure 2.**
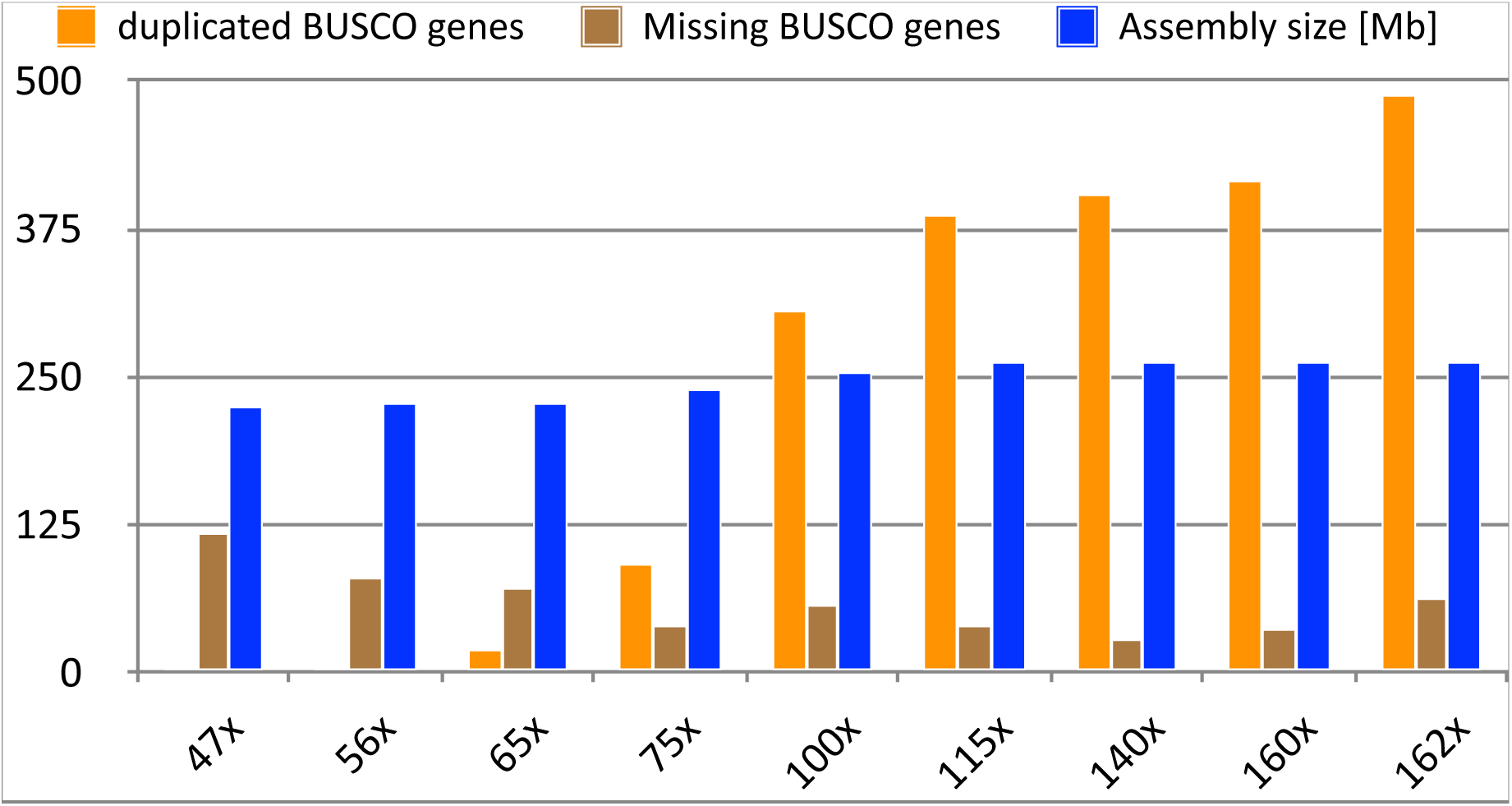
Raw read coverage effect on assembly size and quality. Please note that assembly size is provided in Mb.

By comparing the genome assemblies of relative ants used in the annotation pipeline, our genome assembly seems to have very few N character coverage which means that we have shorter regions between contigs within scaffolds and less portions of input sequencing reads contain N characters (see Table 2). This impact is very much noticeable by comparing assemblies of the closest ants (*P. californicus* and *P. barbatus*) in our set of insects. The N50 of the scaffolds is five times higher because of the about six times higher N character coverage in the *P. barbatus* assembly. Additionally, by considering the assembly size difference of 5 Mb between these two ants we would expect a more complete assembly as well as resulting annotation of the *P. californicus* ant in comparison to the *P. barbatus* ant.

**Table 2.**
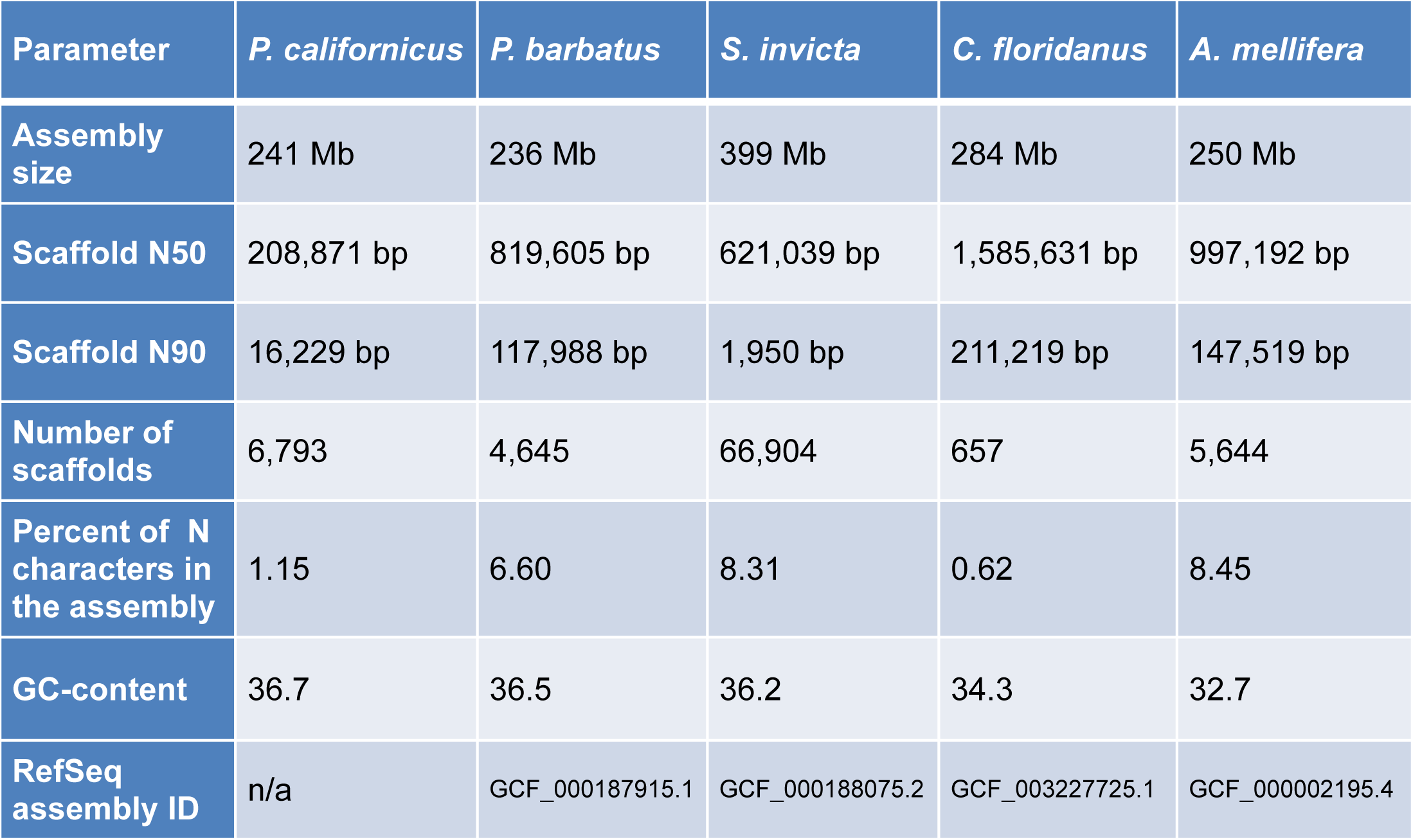
Comparison of genome assemblies of related insect species. With the exception of *P. californicus*, the data were taken from NCBI.

### Annotation of repetitive sequences

Annotation of repetitive sequences was done in two stages. First, we have built a library of repetitive elements, which was later used to annotate individual repeats and mask the genome for annotation of different gene types. We used two different *de novo* pipelines to compile consensus sequences of *P. califormicus* repetitive sequences, namely RepeatModeler (http://repeatmasker.org/RepeatModeler/) and REPET (Flutre et al. 2011). After adding 1,240 Hymenoptera-specific repeats from RepBase, our library consisted of 3,156 sequences, which were subjected to redundancy filtering using CD-HIT with the cut-off level set at 90%. The final library contained 2,595 sequences ranging from 42 to 28,331 bp (median equal to 988 bp). 345 sequences in this dataset were unclassified and TEclass was employed to classify these sequences. We were able to classify most of them and only 71 sequences in our TE library remained unclassified. This library was used as TE-reference set for RepeatMasker run. In total 20.25% of the genome was occupied by repetitive elements, including simple repeats and low complexity regions, 3.98% and 0.53% of the genome, respectively. Not surprisingly, most of the repeats are of TE-origin and all major groups of TEs are represented in the *P. californicus* genome. DNA elements are most common, followed by LTR retroposons and LINEs (see Table 3). Interestingly, SINEs are very rare in the genome. However, it is possible that most of unclassified interspersed repeats are actually SINEs.

**Table 3.** Transposable elements present in *P. californius* genome.

| TE-class | Number of elements | Fraction of the genome |
| --- | --- | --- |
| LTR | 15391 | 4.38% |
| LINE | 9525 | 1.42% |
| SINE | 596 | 0.03% |
| DNA | 72737 | 8.69% |
| Unclassified | 13610 | 1.37% |

### Annotation of protein-coding genes

Homology-based GeMoMa annotation followed by four runs of MAKER2 resulted in 15,688 protein-coding genes, which included 170 exact duplicates of potential transcripts. All following downstream analyses were based on the non-redundant set of 15,518 transcripts and translated proteins of this set referred as being unique. Additionally, 2,288 unique isoforms were annotated by our pipeline based on RNAseq data. Detailed information on number of predicted genes at different stages is provided in Supplementary Figure 1. The missing genes numbers presented in this Figure come from BUSCO assessment on unique *P. californicus* transcripts. Isoforms from the MAKER annotation are referred as alternative transcripts with different intron/exon decomposition (Campbell et al. 2014). For annotation with MAKER, it is recommended to run it at least 3 times. There is a drastic increase of annotation in the second MAKER run, based on the training used SNAP from filtered annotations of the first MAKER run. The high reduction of annotations in MAKER run 3 is based on the training of Augustus on the *P. californicus* genome using BUSCO and forcing detection of start and stop codons in order to predict complete genes.

The functional classification of predicted genes was done employing BLASTp against NCBI’s non-redundant (nr) protein database. We distinguish three categories of functional annotation: a) 8,807 predicted genes were similar to a protein present in nr database with the alignment coverage on query and target of at least 70% of the protein length; b) 3,129 predicted genes were similar to an nr-protein with an alignment coverage of query or target with less than 70% of the protein length, and c) 2047 predicted proteins where neither the query nor the target fulfill the 70% alignment coverage threshold. These predictions show some similarity to proteins but may be novel proteins as they are not clearly classified. 1,535 predicted proteins did not have any cognate protein in nr database. So, in total we classified about 90% of all predicted proteins. These include also 54 proteins that consists of multiple domains potentially representing individual proteins. This may be the result of protein fusion or erroneous gene prediction.

Interestingly, from these 1,535 potential orphan genes from *P. californicus*, 544 are apparently present in the *P. barbatus* genome, however they are missing from current *P. barbatus* annotation. The number of orphan genes or TSG/LSG (taxon specific/lineage specific genes) in *P. californicus* is what would be expected for two relatively closely related ant species but is much lower than what has been shown in the leaf cutter ants (Wissler et al. 2013). In comparison to other relative insects genomes, we have annotated more protein coding genes (see Table 4). This may suggest that our pipeline resulted in some false positive predictions. Interestingly but not surprisingly, non-classified proteins are on average significantly shorter than classified proteins: non-classified proteins are 108 amino acid long on average versus 536 amino acid average length for classified proteins (see Supplementary Figure 2).

**Table 4.** Comparison of *P. californicus genome* annotaVon of related Hymenopteran genomes.

| Species | Protein-coding | tRNA | rRNA | lncRNA | snoRNA | snRNA | Total | Annot. version |
| --- | --- | --- | --- | --- | --- | --- | --- | --- |
| *P. californicus* | 15688 | 1180 | 55 | 931 | 12 | 23 | 17889 | n/a |
| *P. barbatus* | 11348 | 201 | 17 | 1164 | 12 | 19 | 12761 | 101 |
| *S. invicta* | 14473 | 204 | 34 | 1982 | 11 | 26 | 16730 | 103 |
| *C. floridanus* | 11873 | 208 | 30 | 1874 | 13 | 22 | 14020 | 102 |
| *A. mellifera* | 9935 | 218 | 57 | 3146 | 14 | 26 | 13396 | 104 |

In addition to the sequence similarity classification, we also performed further going protein domain analysis with Interproscan. 91% of classified proteins include predictions from Interproscan (see Supplementary Figure 3), while only 24% of non-classified proteins show some Interproscan predictions (see Supplementary Figure 4). The Interproscan results from classified predictions include mostly predictions from PANTHER (Protein ANalysis THrough Evolutionary Relationships), which is a protein classification system (Thomas et al. 2003) and Pfam, which is a large collection of protein domains (El-Gebali et al. 2019). These predictions promote the evidence of the classified proteins. Most predictions of the non-classified proteins are coming from MobiDBLite, which is included in the Interpro database and is used for long intrinsically disordered regions (IDRs) detection (Necci et al. 2017). Based on Intrinsic disorder (ID) and missing Domains in the Pfam database, at least 10% of the human proteome are missing protein domain detections (Mistry et al. 2013). That gives a hint that these proteins are non-classified based on ID and/or incomplete databases. Based on the length of the non-classified proteins (see Supplementary Figure 2), it seems to include several small proteins (SP) which are very much of biological importance but not annotated by most annotation pipelines (Su et al. 2013).

### Odorant receptors

Chemical communication and perception of olfactory cues via ORs is essential for the performance of many tasks in ant colonies (Trible et al. 2017; Yan et al. 2017). Given the biological significance of this gene family in ants, we generated in-depth annotations of OR genes in the two closely related Pogonomyrmex species P. californicus and P. barbatus, for which assembled genomes are available. Our custom pipeline predicted 417 OR gene models in the P. californicus genome and 454 OR gene models in the P. barbatus genome. Of these, 303 gene models were complete in P. californicus and 342 were complete in P. barbatus (see Figure 3, Supplementary Table 2). This nearly doubles the number of originally predicted odorant receptor gene models (274) published for P. barbatus (Smith et al. 2011). Classifying our gene models by known OR gene families showed that most of them fall into the 9-exon (9E) family, with the next biggest families being L, V, E and U in both *P. barbatus* and *P. californicus* (Supplementary Table 3). This is in line with previous studies about OR genes in ants (Engsontia et al. 2015; McKenzie and Kronauer 2018). A phylogenetic analysis of Pogonomyrmex ORs showed that most ORs can be considered as single-copy orthologs, as expected when comparing two closely related species. Clusters in the gene phylogeny of multiple genes from the same species would indicate either very recent gene duplications or losses (i.e. after the species split) or could hint at assembly errors in either genome. The lack of extensive same-species clusters (largest cluster: seven genes, no other cluster exceeding four genes) thus suggests that the assemblies are of equally high quality, with few signs of gene duplication through assembly errors.

**Figure 3.**
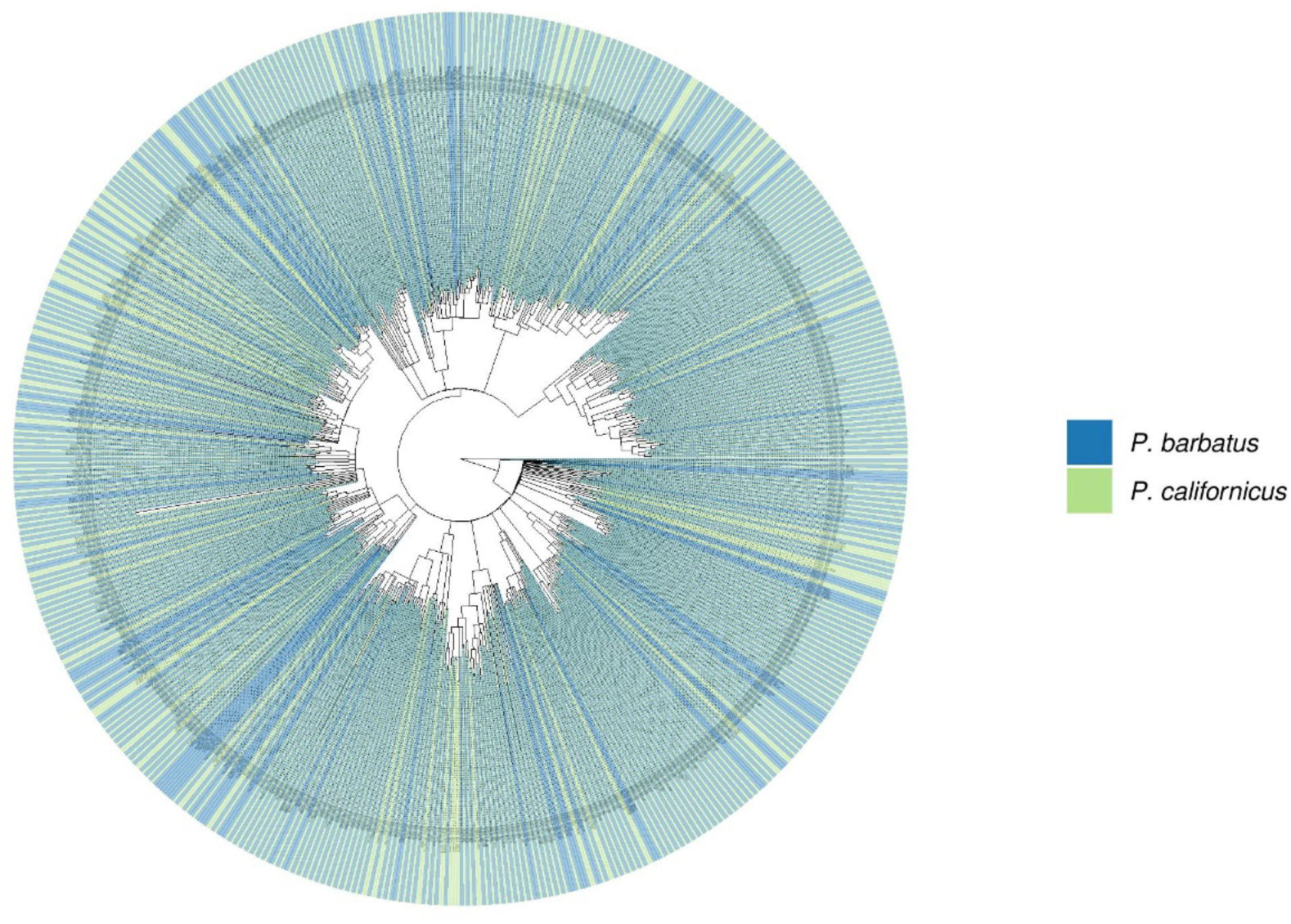
Odorant receptor gene repertoires are similar between P. californicus and P. barbatus. Most gene models have their closest relaVve in the other species. The gene tree shows no large clusters containing genes exclusively of one of the two species. This is evidence for a close relatedness between the species and an equally high quality of the two genome assemblies.

### Annotation of non-protein-coding genes

There are several categories of functional RNAs, including tRNAs, lncRNAs, rRNAs, snoRNAs, snRNAs, and rRNAs. Annotation of some of them is pretty straightforward thanks to the conserved secondary structure, e.g. tRNA or snRNA genes and some are more difficult to annotate, e.g. lncRNA genes. Nevertheless we were able to annotate more than 2,000 such genes in *P. californicus* genome (see Table 4). In general, numbers of non-protein-coding genes detected in *P. californicus* genome are similar to those in other insect genomes with the exception of tRNA genes exceeding more than five times usual number of tRNA genes in insect genomes. Upon close inspection it appeared that the access of tRNA genes is due to unusual number of tRNA^Thr^ genes and in particular its GGT isotype. Moreover, these genes are identical one to each other including 200 bp flanking regions, thus suggesting that they might be an artifact of faulty assembly, not a real biological phenomenon.

### Assembly and annotation quality assessment

BUSCO and DOGMA programs were used for the quality assessment. These programs are working on different signatures in order to estimate the completeness of genome assemblies and resulting annotation of transcripts and proteins. Duplicated transcripts and proteins within annotations of relative genomes were detected using cd-hit as it was done for *P. califonicus* annotation (see Table 5). In general, results from the two programs are in good agreement. The small differences are consequences of different methodology employed by the software; while BUSCO is searching for single-copy orthologous hymenopteran genes, DOGMA searches for Conserved Domain Arrangements (CDA) from an insect reference set. Our annotation of *P. californicus* is comparable or exceeds annotation of published ant genomes. The only parameter that seems to be significantly different in our assembly is a level of genome duplication reported by BUSCO - over 2% comparing to less than 0.4% in other genomes. This is also reflected in number of duplicated transcript but interestingly not in number of duplicated proteins (see Table 5). However, at this point it is difficult to evaluate if this phenomenon reflects intrinsic biological feature of *P. californicus* genome or results from less-than-perfect assembly of the genome.

**Table 5.**
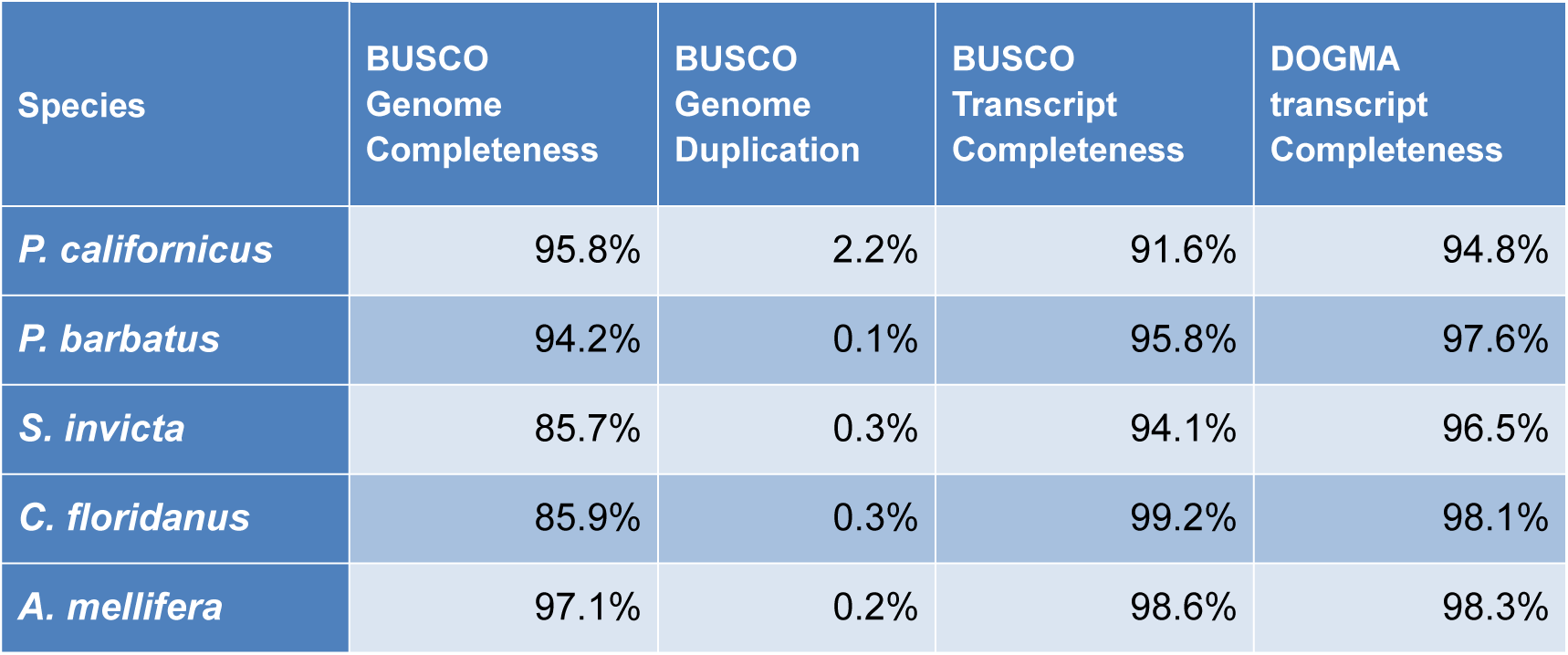
Comparison of completeness and quality of Hymenopteran insects used for the annotation of *P. californicus*. BUSCO and DOGMA analyses are based on the unique sets of transcripts without duplicated sequences and soft masked genomes were used in BUSCO assessments.

## CONCLUSIONS

With the availability of a genome assembly and annotation for *P. californicus* we can now start to analyze the genetic architecture of the intraspecific social polymorphism, differences in aggressive behavior of founding queens, adaptations to desert life in this widely distributed harvester ant. It will also allow us to test whether a supergene, similar to other cases of intraspecific social polymorphism, is responsible for this trait variation. We should also allow to demonstrate that the evolution of OR genes in both *Pogonomyrmex* species proceeded at approximately the same rate without any obvious major gene losses or gains.

## ACKNOWLEDGMENTS

This research was partly funded by the German Research Foundation (DFG) as part of the SFB TRR 212 (NC^3^) – project numbers 316099922 and 396780988 and internal fund of Institute of Bioinformatics.

## Supplementary materials

**Supplementary Figure 1.**
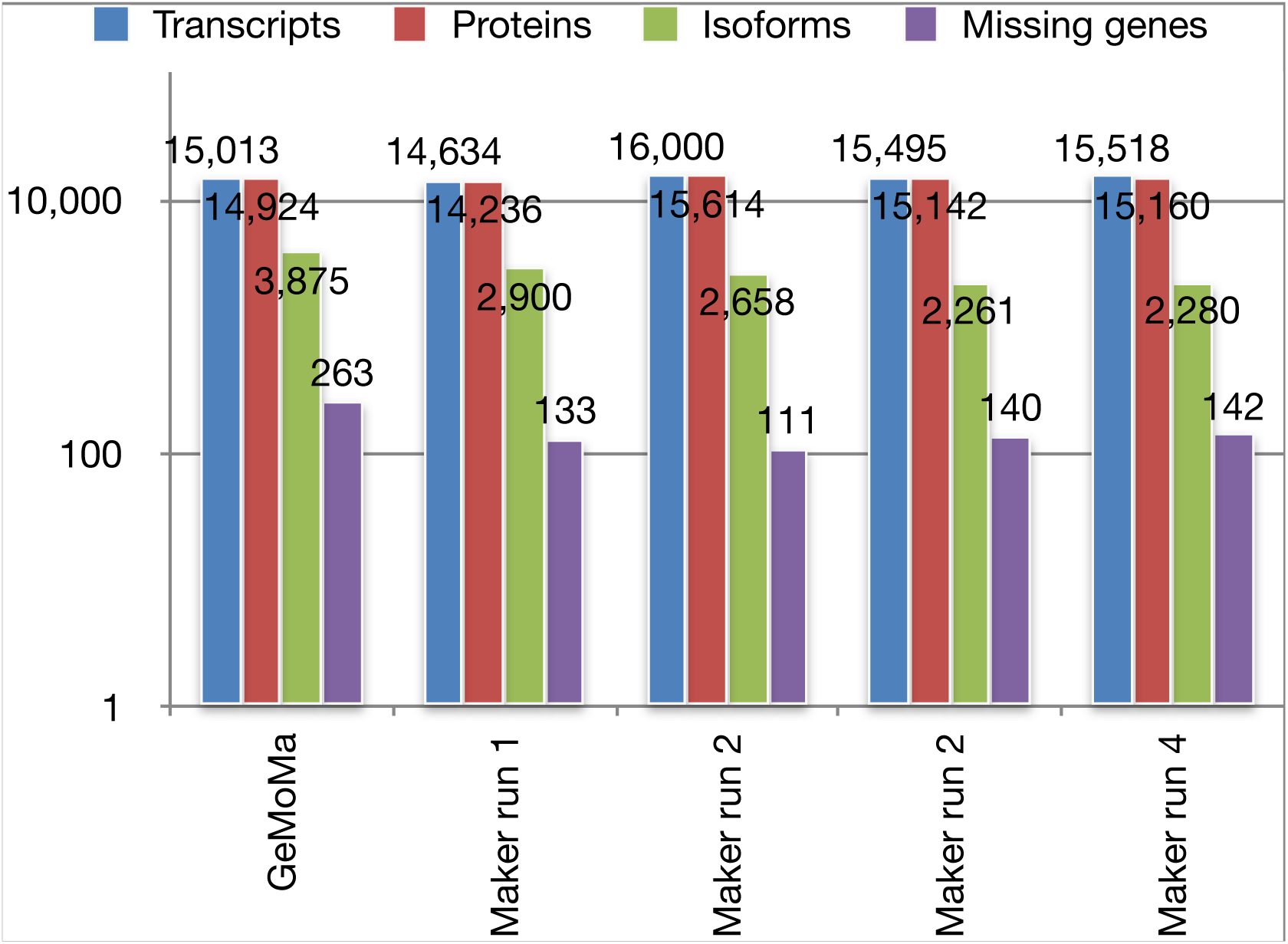
Number of predictions and assessment of completeness through the annotation procedure. This analysis is based on unique/non-redundant sets of transcripts and proteins coming out of the annotation steps.

**Supplementary Figure 2.**
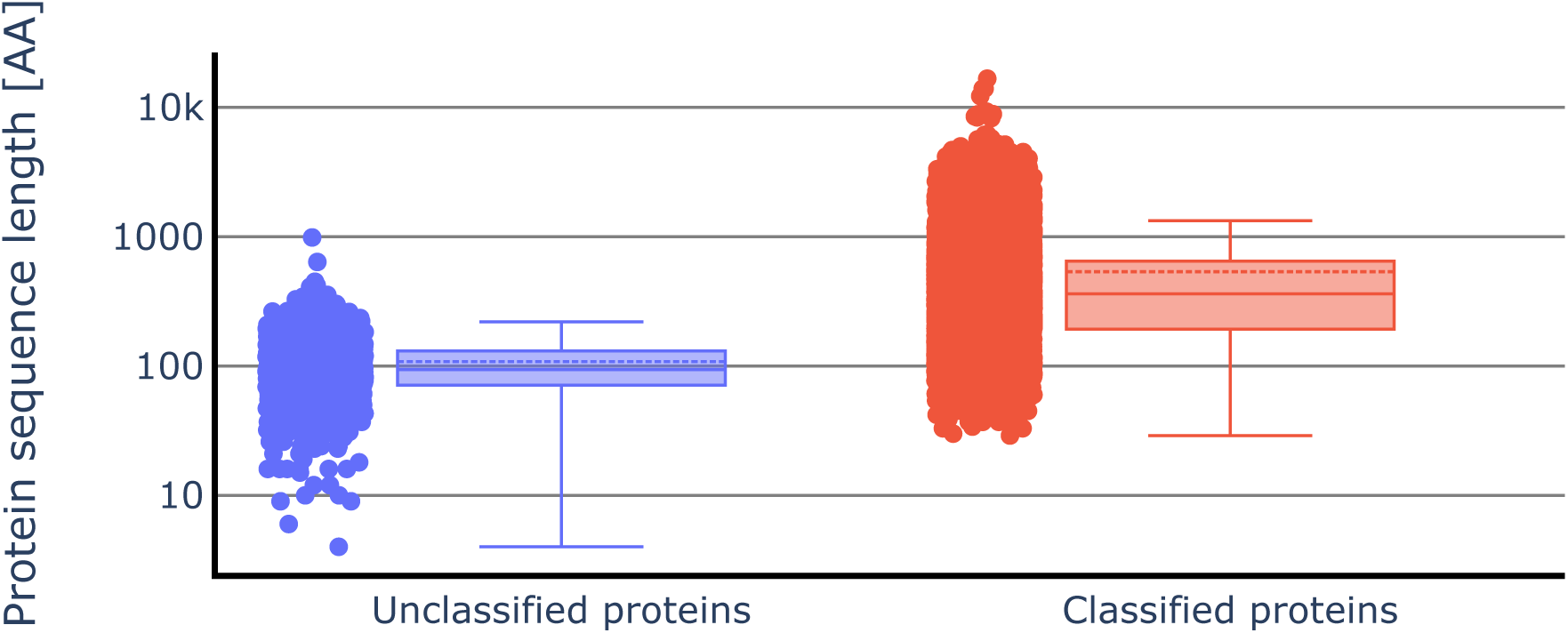
Sequence length distribution of functionally classified and unclassified proteins. Please notice that the y axis is log scaled. The dotted line represents the mean and the solid line the median.

**Supplementary Figure 3.**
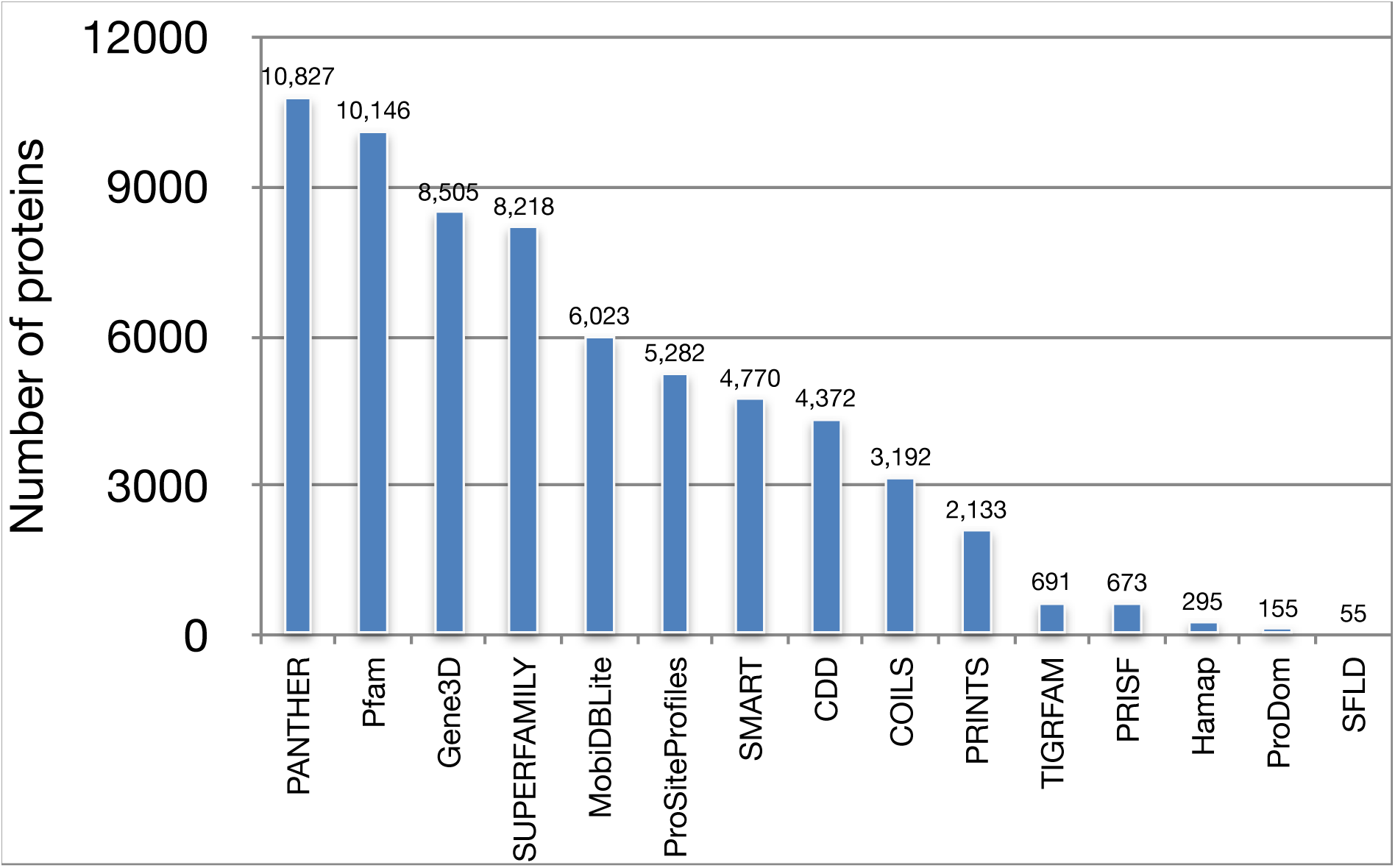
Interproscan prediction distribution of classified proteins.

**Supplementary Figure 4:**
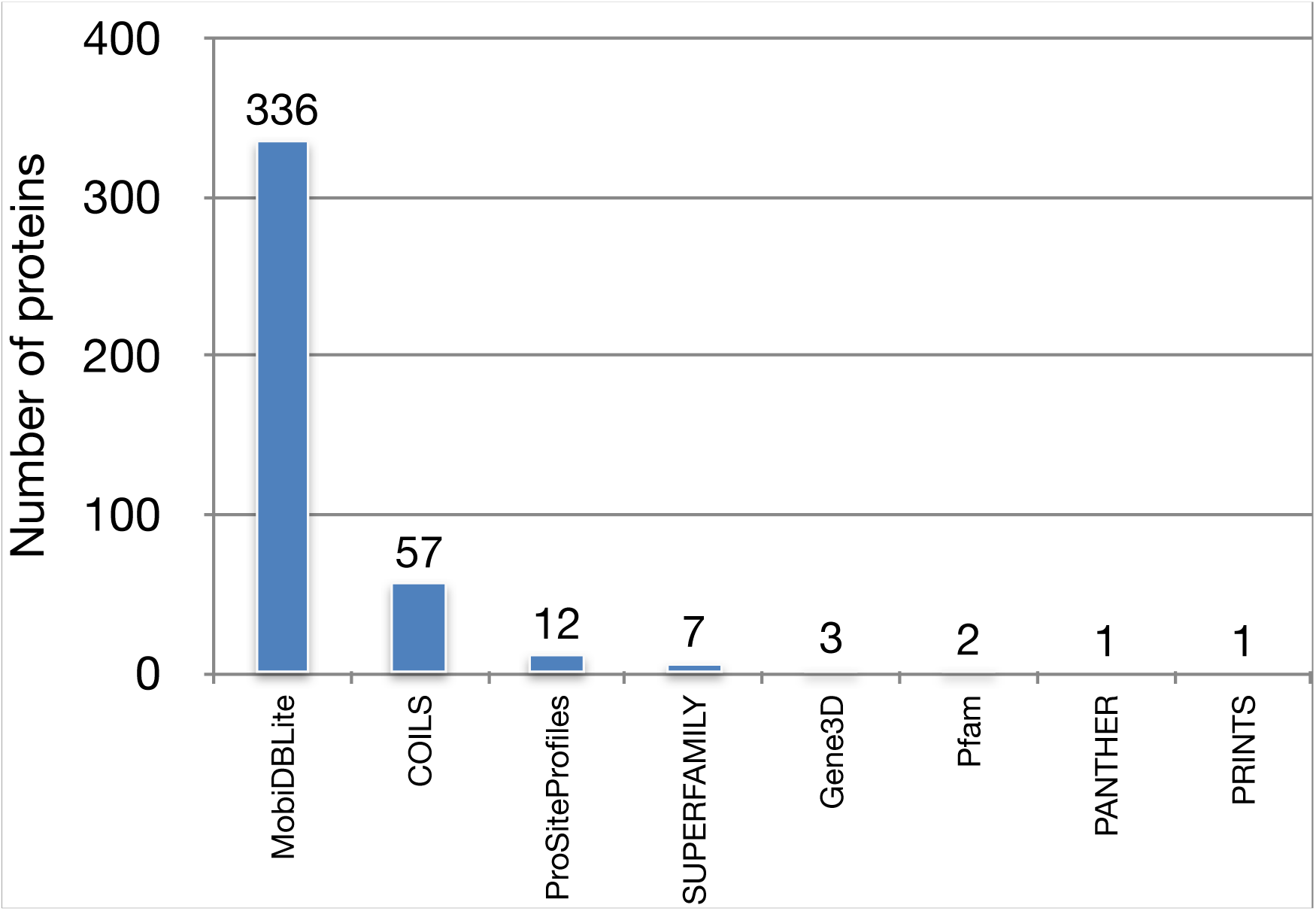
Interproscan prediction distribution of non-classified proteins.

**Supplementary Table 1.** RefSeq accession numbers of assemblies and annotations from relative insects used for the gene annotation.

| Species | RefSeq Accession | Assembly Name | Date of Access |
| --- | --- | --- | --- |
| *P. barbatus* | GCF_000187915.1 | Pbar_UMD_V03 | May 2018 |
| *S. invicta* | GCF_000188075.1 | Si_gnH | May 2018 |
| *C. floridanus* | GCF_003227725.1 | Cflo_v7.5 | July 2018 |
| *A. mellifera* | GCF_000002195.4 | Amel_4.5 | May 2018 |

**Supplementary Table 2.** Number of OR gene models in the annotation of the *P. barbatus* genome available in NCBI as well as those predicted by the custom pipeline for both the *P. californicus* and *P. barbatus* genomes. Incomplete gene models are either missing N- and/or C-termini or, in case of the original annotation of *P. barbatus*, could not be identified as OR genes, but contain an OR domain. Homology based predictions do not contain an OR domain at all.

| Species assembly | All OR gene models | Perfect OR gene models | Incomplete OR gene models | Homology based OR gene models |
| --- | --- | --- | --- | --- |
| *P. barbatus* (manual annotation) | 274 | 109 | 165 | 0 |
| *P. californicus* | 417 | 303 | 67 | 47 |
| *P. barbatus* | 454 | 342 | 43 | 69 |

**Supplementary Table 3.** Number of complete, incomplete, and homology based gene models predicted by the custom pipeline for both *P. californicus* and *P. barbatus* in the OR gene families.

| OR gene family | Pcal complete | Pcal incomplete | Pcal domain missing | Pbar complete | Pbar incomplete | Pbar domain missing |
| --- | --- | --- | --- | --- | --- | --- |
| 9E | 118 | 32 | 23 | 141 | 22 | 37 |
| A | 9 | 1 | 3 | 10 | 1 | 2 |
| B | - | 1 | - | 1 | - | - |
| C | - | 1 | 1 | 1 | - | - |
| D | 3 | - | - | 4 | - | - |
| E | 24 | 5 | 3 | 28 | 2 | 6 |
| F | 8 | - | - | 8 | - | - |
| G | 1 | 1 | 6 | 2 | - | 9 |
| H | 10 | 6 | 3 | 12 | 1 | 1 |
| I | 1 | - | - | 1 | - | - |
| J | 3 | 1 | - | 2 | - | - |
| K | 2 | - | - | 2 | - | - |
| L | 42 | 3 | 1 | 44 | - | 4 |
| M | 4 | - | - | 4 | - | - |
| N | 4 | - | - | 4 | - | - |
| Orco | 1 | - | - | 1 | - | - |
| P | 9 | - | - | 8 | - | - |
| Q | 1 | - | - | 1 | - | - |
| R | 1 | 1 | - | 1 | - | 1 |
| S | 1 | - | - | 1 | - | - |
| T | 1 | 3 | - | 6 | - | - |
| U | 18 | 7 | 1 | 19 | 13 | 3 |
| V | 40 | 5 | 3 | 39 | 4 | 4 |
| W | 1 | - | - | 1 | - | - |
| X | - | - | 1 | - | - | 1 |
| XA | - | - | - | - | - | 1 |
| Z | 1 | - | 2 | 1 | - | - |

